# Evolving Changes in Cortical and Sub-Cortical Excitability during Movement Preparation: A Study of Brain Potentials and Eye-Blink Reflexes during Loud Acoustic Stimulation

**DOI:** 10.1101/2020.08.19.258327

**Authors:** An T. Nguyen, James R. Tresilian, Ottmar V. Lipp, Dayse Tavora-Vieira, Welber Marinovic

## Abstract

The presentation of Loud Acoustic Stimuli (LAS) during preparation can trigger motor actions at very short latencies in a phenomenon called the StartReact effect. It was initially proposed that a special, separate sub-cortical mechanism which by-passes slower cortical processes could be involved. We sought to examine the evidence for a separate mechanism against the alternative that responses to LAS can be explained by a combination of stimulus intensity effects and preparatory-states – as proposed by activation models of motor control.

To investigate whether cortically mediated preparatory processes are involved in shaping reactions to LAS, we used an auditory reaction task where we manipulated preparation-level within each trial. We contrasted responses to non-intense tones and LAS and examined whether cortical activation, sub-cortical excitability (measured by pre-stimulus EEG and eye-blink startle reflexes, respectively) and the motor response were influenced by preparation-level.

As predicted by the activation model, increases in preparation-level were marked by gradual reductions in RT coupled with increases in cortical activation and sub-cortical excitability – at both condition- and trial-levels. Changes in cortical activation influenced motor and auditory but not visual areas – highlighting the wide-spread yet selective nature of preparation. RTs were shorter to LAS than tones, but the overall pattern of preparation-level effects were the same for both stimuli. These results demonstrate that LAS responses are indeed shaped by cortically mediated preparatory processes. The concurrent changes observed in brain and behaviour with increasing preparation reinforces the notion that preparation is marked by evolving brain states which shape the motor response.

**Key Points:**

- Reactions to Loud Acoustic Stimuli can be explained by stimulus intensity and preparation state
- We manipulated movement preparation by altering the temporal position of the imperative stimulus
- Preparation was marked by reductions in RT, and increased cortical and sub-cortical excitability
- Preparation had the same effect on reactions to Loud Acoustic Stimuli and non-intense tones
- The results highlight the widespread, evolving, and strategic nature of movement preparation

## Introduction

Before the execution of a volitional movement, neural processes prepare the neuromotor system to perform that action (Requin et al., 1991; Requin & Riehle, 1995). In reaction time (RT) tasks and anticipatory timing tasks, these processes establish a state of readiness to respond to a stimulus (R. Chen et al., 1998; Ibáñez et al., 2020; Schmidt et al., 2018). This readiness is marked by increased activity in response-related circuits both within the brain and the spinal cord (R. Chen et al., 1998; Eichenberger & Rüegg, 1984; Leocani et al., 2000). This increased activity is associated with increased sub-cortical excitability, such as stretch reflex excitability in response-related musculature (Sullivan & Hayes, 1987), as well as increased cortical excitability in the circuitry responsible for generating the volitional response (Hoffstaedter et al., 2013; Toro et al., 1994).

In RT and anticipatory timing tasks, prepared actions can be triggered at very low latencies by the presentation of an intense stimulus, such as a loud acoustic stimulus (LAS): the StartReact effect. It was initially theorised that these responses were mediated by a separate mechanism that by-passes the cortex, involving sub-cortical circuitry associated with the startle response (Carlsen et al., 2004; Valls-Solé et al., 1999). This proposal was based on the idea that mental representations which specify the parameters of the movement (motor programmes) are prepared in advance and transferred to sub-cortical structures for storage and execution (Carlsen et al., 2004; Valls-Solé et al., 1999). The presentation of the intense stimulus was thought to excite these structures as part of the startle response, involuntarily triggering the release of the motor programme while by-passing slower cortical triggering mechanisms involved in voluntary motor control.

Recent studies suggest that the cortex may not be by-passed in the StartReact effect as initially suggested (Alibiglou & MacKinnon, 2012; Marinovic et al., 2014; Stevenson et al., 2014). This idea aligns with our proposal (see Marinovic & Tresilian, 2016) that this phenomenon is mediated by voluntary control pathways and could be the result of stimulus intensity effects (Cattell, 1886; Piéron, 1913). If this were the case, the dynamic changes that occur cortically during movement preparation would be expected to drive the manifestation of the StartReact effect. In activation models (See Figure 1), changes in excitability during movement preparation are visualised as the build-up of activation over time as the expected moment of response initiation approaches, and responses are initiated when activation reaches a certain threshold (Tresilian & Plooy, 2006). Stimulus-evoked activity is also modelled to increase activation in an additive manner as suggested by McInnes et al. (2020), proportional to the intensity of the stimulus. Responses are modelled to be shaped by a combination of these factors, where higher levels of stimulus intensity (Figure 1A) and preparation (Figure 1B) are associated with shorter RTs.

**Figure 1.**
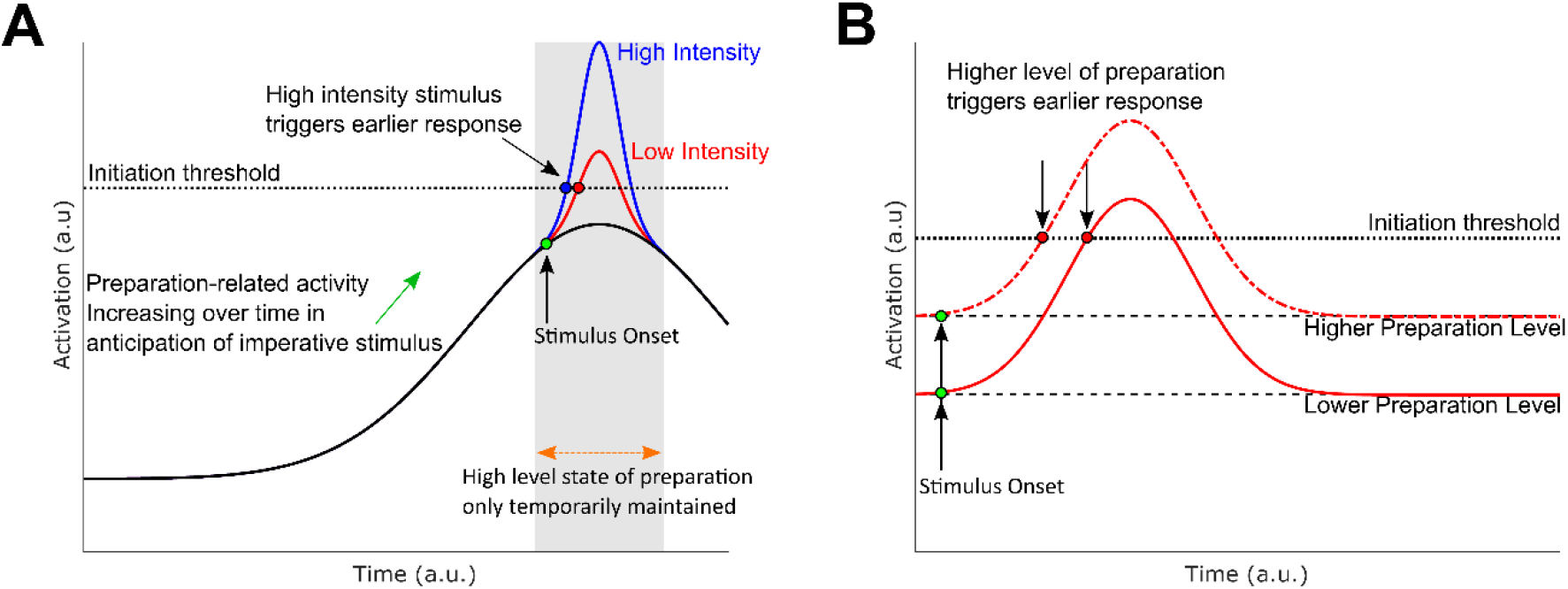
(**A**) Conceptual visualisation of the Activation Model, depicting activation in the motor system from the start of the trial, leading up to a response. The black line represents preparation-related activity which gradually increases as expected time of the response approaches. The grey area shows that a high level of preparation can only be maintained for a short period of time. The red and blue lines represent the activity evoked by low and high intensity acoustic stimuli, respectively. This induced activity causes the net activity in the system to cross the initiation threshold (dotted black line), triggering the response; but activity in the high intensity stimulus reaches the threshold earlier, producing an earlier response. (**B**) Visualisation showing the influence of preparation level on response time, given the same stimulus (static levels of preparation used for simplicity). When the system is at a higher state of preparation, voluntary responses to stimuli can occur earlier because less additional activation is required to reach initiation threshold.

**Figure 1.**
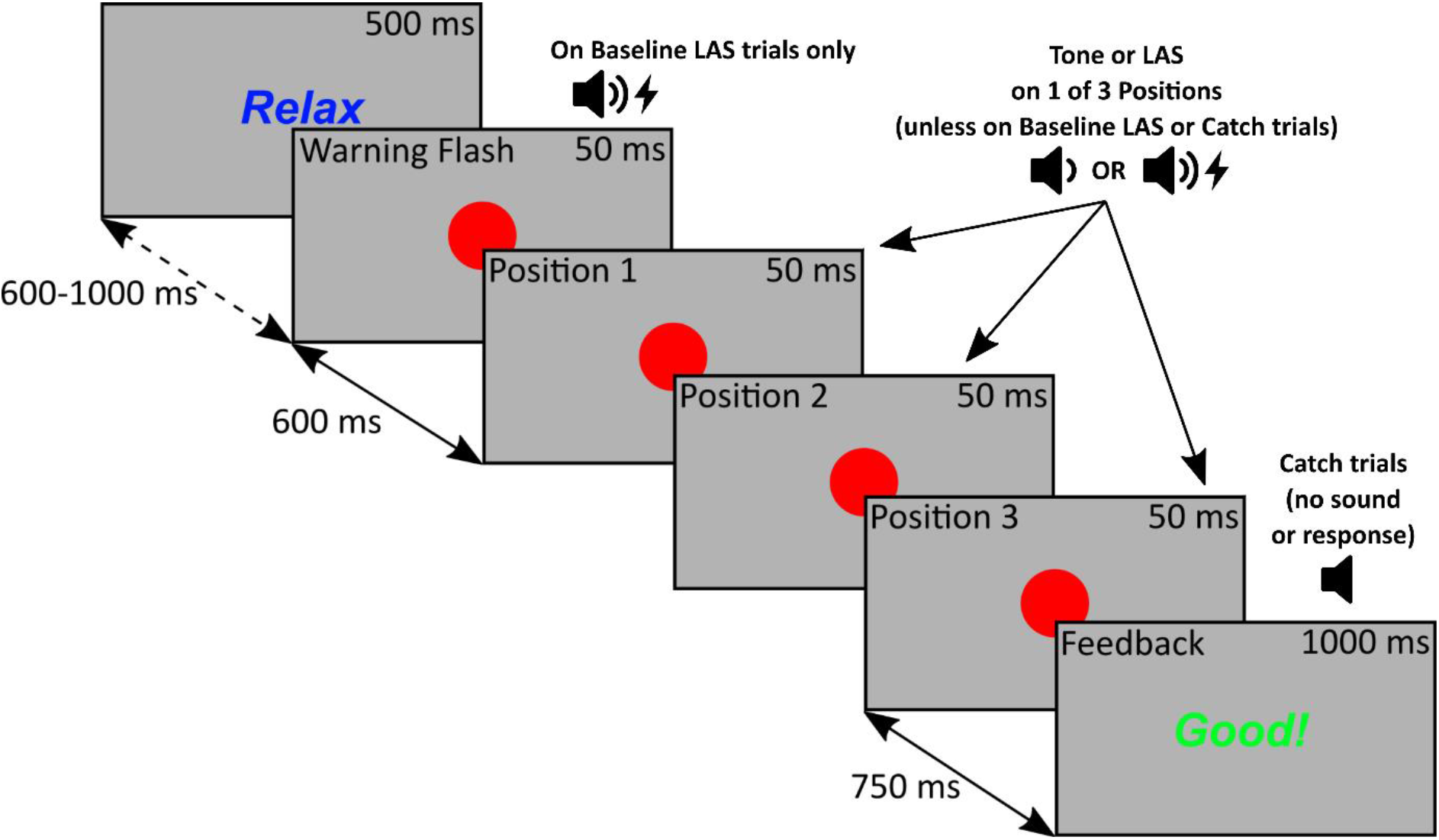
A diagram showing the sequence of events on each trial. A red circle was briefly flashed on-screen four times and an acoustic stimulus, either the tone or loud acoustic stimulus (LAS) was randomly presented with the flash at Positions 1, 2 or 3. Participants reacted to the sound by pressing on a force sensor, and feedback about timing and force was presented at the end of the trial. On a small portion of trials, the LAS was presented with the Warning Flash (Baseline LAS trial), or no sound is presented at all (Catch trials) - No responses were required on these trials.

In this model, variability in RTs across trials are attributed to fluctuations in motor preparation, in part due to uncertainties about precisely when responses will be required. As high levels of preparation can only be maintained for a short duration (Alegria, 1974; Müller-Gethmann et al., 2003), accurate responding relies on the appropriate timing of preparatory-related processes which are informed by previous knowledge and experience, and current information. Preparation is relatively straightforward in predictable tasks (e.g., anticipatory timing tasks, and RT tasks with a fixed foreperiod) leading to overall shorter RTs. However, it is more difficult to maintain a high level of preparation in tasks with temporal uncertainty (e.g., tasks with a random foreperiod), leading to relatively longer and more varied RTs (Leow et al., 2018).

Consistent with this account, it has recently been shown that higher levels of temporal preparation were associated with graded decreases in RT to both non-intense tones and LAS in an unpredictable reaction task (Leow et al., 2018). Concerning cortical and sub-cortical excitability, studies have used LAS to probe the time-course effects of movement preparation separately for each system. For example, cortical preparation-related activity reflected by the contingent negative variation (CNV) – a slow and centrally distributed negativity in the scalp electroencephalogram (EEG) implicated in the anticipation for an upcoming stimulus and movement preparation (Kononowicz & Penney, 2016) – was associated with RT in a predictable RT task (MacKinnon et al., 2013), and some have reported that reflex excitability is enhanced around the expected time of the response compared to baseline (Carlsen et al., 2004; Lipp et al., 2001; Marinovic et al., 2013; Valls-Solé et al., 1999; Valls-Solé et al., 1995). However, research has yet to systematically examine how both cortical and sub-cortical excitability are influenced by the level of movement preparation, and whether they, in turn, are associated with the motor action.

### Current Study

In this study, we sought to capture the evolution of cortical and sub-cortical excitability that occurs over time with motor-preparation and examine its relationship with voluntary responses to LAS and non-intense stimuli. To study the evolving effects of motor preparation, we modified an auditory RT task to induce increasing levels of motor preparation within each trial. We expected that RTs to LAS would be significantly shorter than non-intense tones, reflecting an effect of stimulus intensity. According to the activation model, responses to LAS and non-intense tones are expected to be similarly influenced by the level of preparation – such that RTs should decrease, and forces should increase as the sounds are presented in later positions, demonstrating that increased readiness to respond leads to faster responses.

Regarding cortical excitability, we expected to observe an increasing negativity in the motor region as sounds are presented in later positions, reflecting increasing levels of preparation. This pattern should be observed on both tone and LAS trials, demonstrating cortical involvement in both contexts. Extending beyond the motor system, recent research has shown interactions between auditory and motor regions of the brain during speech and musical rhythm perception in humans (J. L. Chen et al., 2008; J. L. Chen et al., 2006; Cheung et al., 2016). Work by Li and colleagues (2017) has further suggested that the auditory cortex is required not only to anticipate a sensory event but also produce appropriate motor responses in mice. As such, we also sought examine whether activity in sensory cortical areas (auditory and visual) would also evolve with preparation. Lastly, we also expected that sub-cortical excitability would increase as the level of motor preparation evolves over time, reflected by decreases in blink latency and increases in blink amplitude.

## Method

### Participants

Thirty-one healthy adult participants consisting of university student and staff volunteers were recruited. Eight participants were excluded in total; two due to excessive EEG noise and artifacts, two due to low performance on ‘Catch’ trials (< 70%), and four due to missing behavioural data. The final sample consisted of 23 participants (age *M*(*SD*) = 20.43(2.57) years, age range = 18 – 27 years, 18 females). All participants reported being right-hand dominant, having normal or corrected to normal vision, no history of significant head trauma and no diagnosed neurological conditions. All participants provided written informed consent before starting the experiment and the protocol was approved by the human research ethics committee of Curtin University (approval code: HRE2018-0257).

### Modified Auditory Reaction Task

Participants were instructed to quickly respond to an auditory stimulus (tone or LAS) that was randomly embedded in a sequence of four visual flashes (See Figure 1). This design allowed us to manipulate the level of motor preparation, which can be represented by the conditional probability (See Table 1). The tone was a 1700 Hz pure tone presented for 50 ms at 60 dBa, and the LAS was a broadband white-noise stimulus presented for 50 ms at 104 dBa. Participants responded by pressing on a force sensor (SingleTact, Model: CS8-10N) with their right index finger. The force sensor was embedded in a custom-built device resembling a computer mouse. The task was presented using MATLAB 2015b and Psychtoolbox version 3.0.11 (Brainard, 1997; Kleiner et al., 2007; Pelli, 1997) on an ASUS 24-inch LCD monitor (Model: VG248QE, running at 1920 x 1080 resolution and 120 Hz).

**Table 1.** Summary showing the number of tone and loud acoustic stimulus (LAS) trials for each Position against the number of possible remaining events. Note the evolving chance of hearing an auditory stimulus at each position of the task presented on the bottom row.

| Stimulus Type | Warning Flash | Position 1 | Position 2 | Position 3 |
| --- | --- | --- | --- | --- |
| Tone | 0/360 | 80/340 | 80/240 | 80/140 |
| LAS | 20/360 | 20/340 | 20/240 | 20/140 |
| Total | 20/360 | 100/340 | 100/240 | 100/140 |

| Conditional probability | 5.55% | 29.41% | 41.67% | 71.42% |
| --- | --- | --- | --- | --- |

The auditory stimuli were presented through Sennheiser headphones (Model: HD25-1 II). The recorded rise time of both stimuli from the output of a soundwave was <1.25 ms. Sound intensity was measured with a Brüel & Kjaer sound-level meter (type 2205, A weighted) placed 2 cm from the headphone speaker.

On each trial, ‘Relax’ was presented for 500 ms followed by a blank screen ranging randomly from 600 – 1000 ms. A red circle (42 mm in diameter) was briefly flashed on the centre of the display four times (50 ms duration with a stimulus onset asynchrony of 600 ms). The first flash served as the warning cue while the following three flashes served as potential positions where the tone could be presented (referred to as Positions 1, 2 and 3). Occasionally, a LAS was presented instead of the tone. Visual feedback was presented 750 ms after the final flash for 1000 ms. On some trials, the LAS was paired with the warning flash to elicit a baseline measure of the eye-blink startle reflex. To discourage anticipatory responding to the final flash, we also included Catch trials where no auditory stimulus was presented, and no response was required. The order of trials was pseudo-randomised such that the LAS was never presented on two consecutive trials.

The experimental portion of the task consisted of 360 trials, split into 4 blocks of 90 trials with self-paced breaks between blocks. In total, there were 240 tone trials (66.67% of total trials, 80 trials for each Position), 80 LAS trials (22.22% of total trials, 20 for each Position, including the warning flash), and 40 Catch trials (11.11% of total trials). Before the experimental task, participants were provided with verbal and on-screen instructions, example demonstrations of the trial sequence, the tone and LAS, followed by a practice block consisting of 18 trials in a fixed sequence with the same trial proportions as the experimental blocks.

With respect to feedback, ‘Good Timing’ was presented on tone trials if participants responded between 50 - 250 ms after stimulus-onset. Otherwise, ‘Too early’ or ‘Too late’ was presented. ‘No response detected’ was presented if no response was made within 600 ms. On Catch trials, ‘Good’ was presented if no response was detected, otherwise ‘Oops!’ was presented. On LAS trials, no feedback on performance was presented, and ‘Probe trial’ was presented in place of feedback. A point system was also implemented to encourage task engagement. Three points were awarded for timely responses on tone trials, but points were not deducted for inaccurate or absent responses. No points were awarded or deducted for Catch or LAS trials, and points were reset after each block. Points were not recorded or analysed.

### Force and EMG Data Reduction and Measurement

Force data were continuously recorded for the duration of the trial, digitised at 2000 Hz using a National Instruments data acquisition device (Model: USB-6229). The data were filtered using a low-pass second-order Butterworth filter with a cut-off frequency of 20 Hz. We measured movement onset, relative to the onset of the auditory stimulus, calculated from the tangential speed time-series derived from the force data using the algorithm recommended by Teasdale and colleagues (1993). Trials with response times outside 50 - 600 ms were excluded from further analysis, *M(SD)* = 10.95(12.31) trials (~2%). We also measured the peak force of each response.

We recorded EMG activity from the right Orbicularis Oculi muscle using Ag/AgCl sintered electrodes in a pre-amplified bi-polar set-up. One electrode was placed below the pupil, the second was placed laterally and slightly higher than the first electrode, ~1 cm edge-to-edge. A ground electrode was placed on the right mastoid region. We used a Neurolog Systems Digitimer Pre-Amplifier (Model: NL820) and Amplifier (Model: NL905), with a 50 – 1000 Hz pass-band filter and Gain set to 1000. The data were also digitised using the National Instruments DAQ.

The EMG data were processed offline using a semi-automated procedure in R. Firstly, the data were down-sampled to 1000 Hz, rectified using the ‘rectification’ function in ‘biosignalEMG’ package (Guerrero & Macias-Diaz, 2018). The Bonato et al. (1998) method was used to automatically detect blink onset latency on the rectified data, using the ‘onoff_bonato’ function in the ‘biosignalEMG package’ (sigma n = standard deviation of activity 0 – 200 ms prior to the LAS). If no onset was detected, the threshold was gradually increased (up to 3 times sigma n) then decreased (down to 0.5 times signa n), until an onset was detected within 20 – 150 ms.

We measured baseline-to-peak EMG amplitude occurring after blink onset on the smoothed data using a 5-point moving average with the ‘rollapply’ function in the ‘zoo’ package (Zeileis et al., 2020). Each trial was manually screened, and corrections were made to onset and peak latencies where necessary. Acceptable onset latencies were within 20 to 80 ms from LAS onset, trials outside this window were excluded from further analyses of blink data (Blumenthal et al., 2005). Trials with a flat EMG response were classified as ‘non-response trials’, trials containing excessive noise, artifacts, or voluntary activation before 20 ms were classified as ‘missing’ trials. Non-response and missing trials were not included in further analyses of blink data. On average, 3.35(4.49) trials (~4.2%) per participant were identified as non-response, 3.35(4.91) trials (~4.2%) were identified missing, 5.96(6.06) trials (~7.5%) were manually adjusted.

### EEG Data Acquisition, Pre-Processing and ERP measurement

EEG data were recorded continuously throughout the experimental blocks. Data were acquired using a Biosemi ActiveTwo EEG system and ActiView (ver. 7.07) at a sampling rate of 2048 Hz with a 100 Hz low-pass online filter. Data were recorded from 64 scalp electrodes arranged according to the 10-5 system with additional electrodes placed adjacent to the outer canthi of the left eye and on the left infraorbital region. For online referencing, the Biosemi EEG system uses active electrodes with Common Mode Sense and Driven Right Leg electrodes providing a reference relative to the amplifier reference voltage.

EEG data were processed offline in MATLAB 2018a using EEGLAB (Delorme & Makeig, 2004), AMICA (Palmer et al., 2012), SASICA (Chaumon et al., 2015), and ERPLAB (Lopez-Calderon & Luck, 2014) plugins. The data were re-referenced to the average of the 64 scalp electrodes, filtered from 0.1 – 40 Hz with separate low- and high-pass filters, using the ‘pop_eegfiltnew’ function in EEGLAB. The filtered data were then down sampled to 256 Hz.

Epochs were extracted on tone and LAS trials, time-locked to the onset of the sound. Epochs spanned for the entire trial (−5000 to 3000 ms) and baseline amplitudes were corrected to the 100 ms interval before the previous flash (i.e., −700 to −600 ms relative to the sound). A close baseline was chosen to minimise the influence of different foreperiod length on amplitude measures, allowing us to focus on changes in expectation from one Position to the next. Figure 2 shows ERP waveforms which are baseline-corrected to the start of the trial, whereas the ERP waveforms in Figure 3 show data baseline-corrected to the previous flash. To investigate the impact of different baselines, we contrasted results between pre-warning cue and previous-flash baselines. Pre-warning baselined led to larger differences across Position conditions on amplitude but did not have a meaningful impact on the overall pattern of results. To correct for blinks, horizontal saccades and other artifacts, Independent Component Analysis was conducted and independent components (ICs) containing artifacts were manually identified with the guidance of SASICA and removed, *M*(*SD*) = 12.3(5.6) ICs.

**Figure 2.**
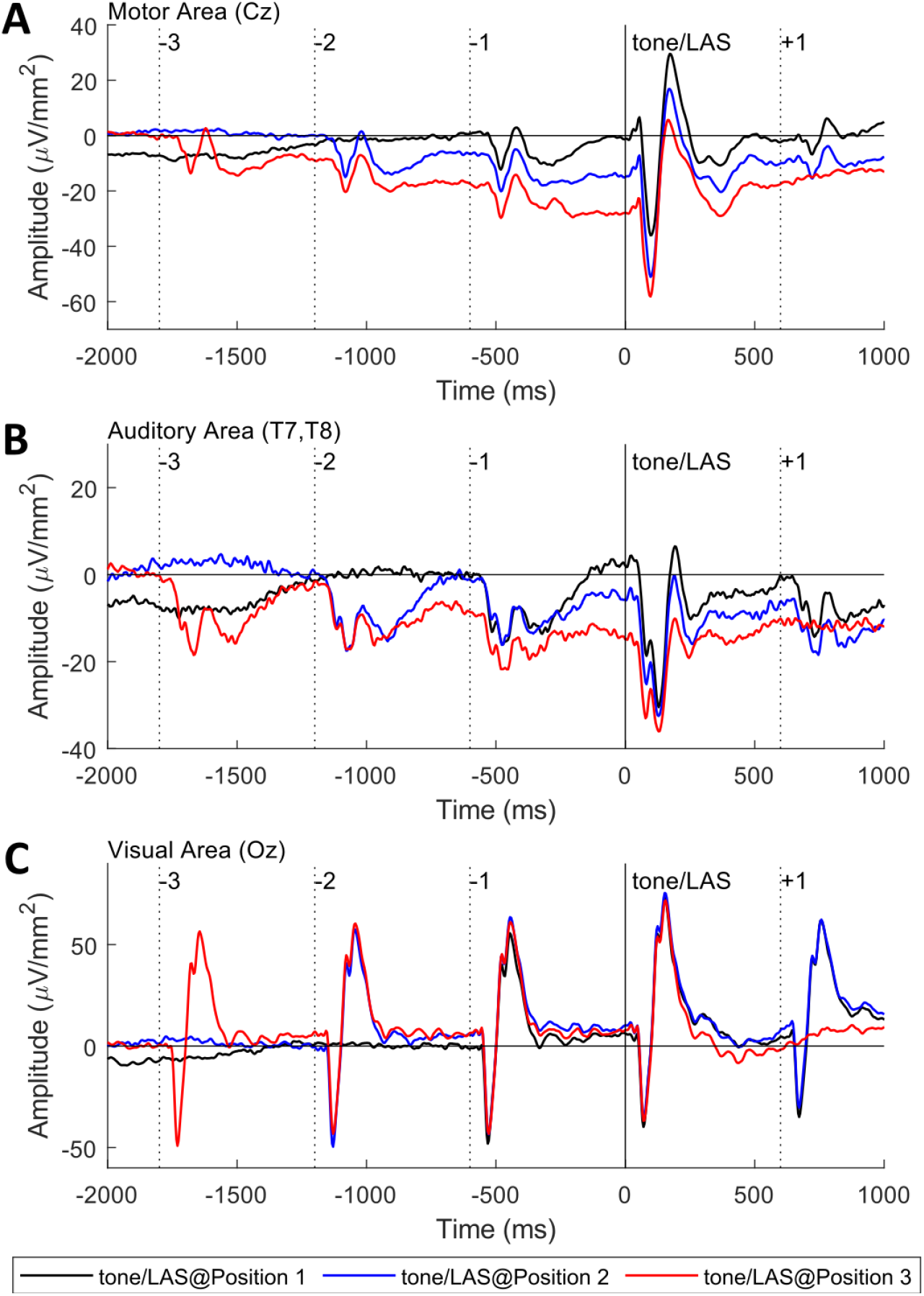
Grand-averaged ERP waveforms at motor (**A**), auditory (**B**) and visual areas (**C**) for each position (1,2,3), averaged across stimuli (tones, LAS). Plots are aligned to the onset of the imperative stimulus, and are baseline corrected to −100 to 0 ms prior to the warning cue. The figure shows the evolution of cortical activation over the course of the trial, which is evident in motor and auditory, but not visual areas.

**Figure 3.**
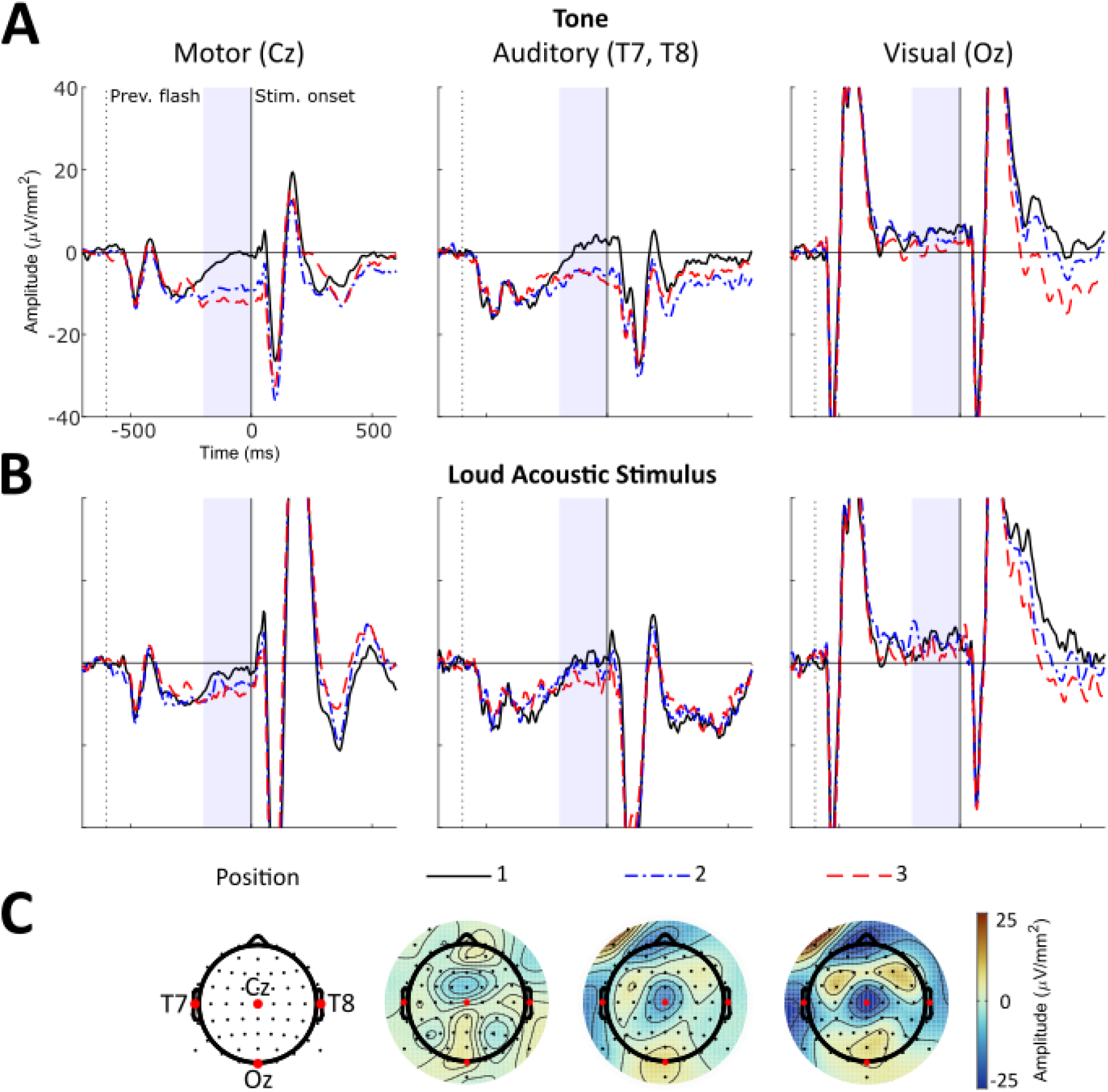
Grand-averaged waveforms at each Position for tone (**A**) and loud acoustic stimulus trials (**B**), at scalp sites corresponding to motor (Cz), auditory (T7, T8) and visual areas (Oz). Waveforms are aligned to the onset of the auditory stimulus and baseline-corrected to −100 to 0 ms relative to the previous flash. The blue shaded area shows the interval where pre-stimulus mean amplitudes were measured. (**C**) Topographical maps show the distribution of activity during the measured interval on tone trials.

**Figure 3.**
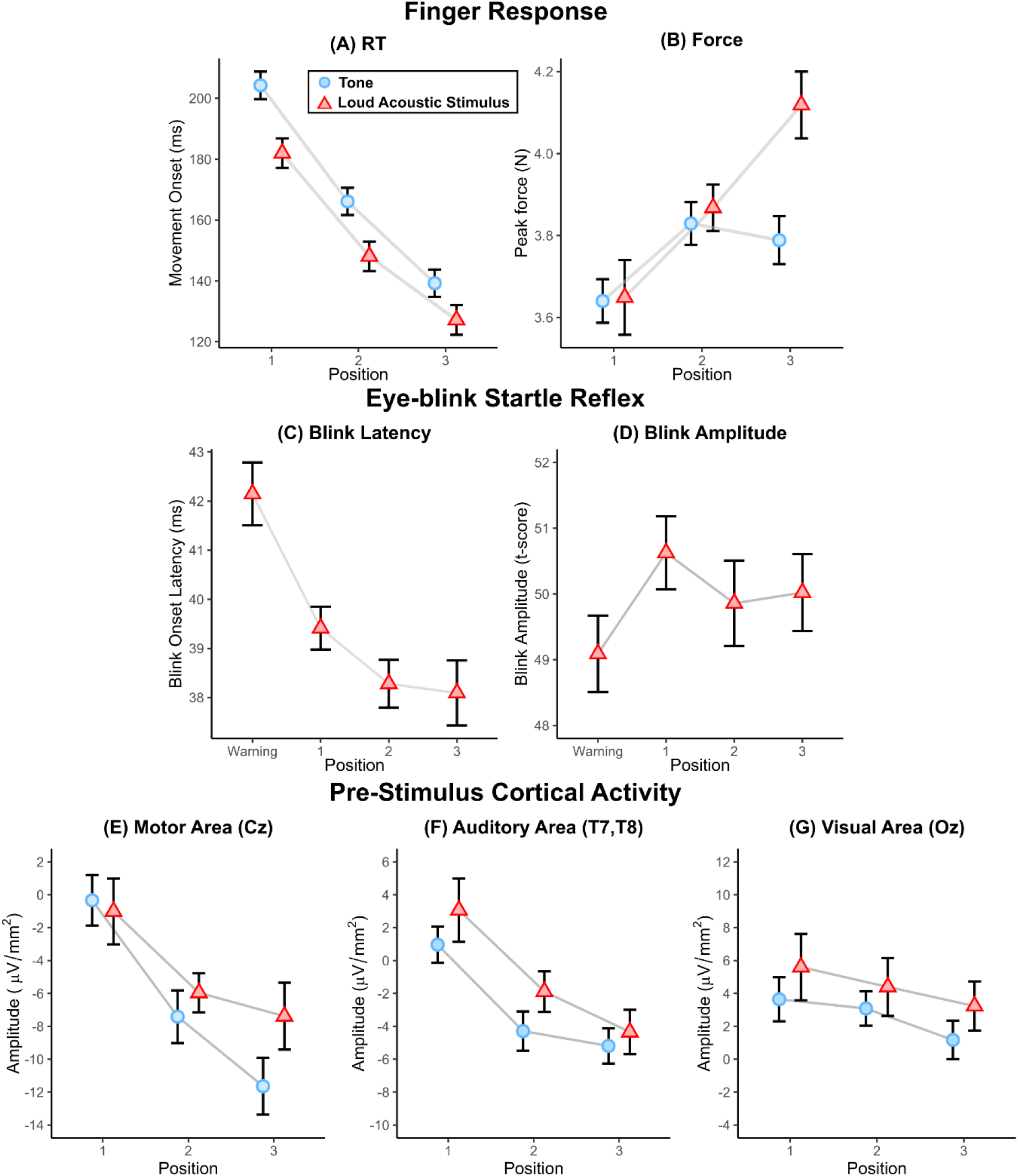
Grand means and within-subject standard error bars for **(A)** RT, **(B)** force, **(C)** blink onset latency, **(D)** blink amplitude, and pre-stimulus cortical activity at **(E)**motor, **(F)** auditory and **(G)** visual areas.

A Surface Laplacian filter was applied using algorithms described in Perrin and colleagues (1989) (smoothing factor = 1e^−5^, order of Legendre polynomial = 10) to reduce volume conduction effects in EEG sensor space, resulting in a μV/mm^2^ voltage scale. Trials containing voltages on analysed channels exceeding ± 100 μV/mm^2^ were excluded, *M*(*SD*) = 8.45(7.40) trials (~2.3%). After trial rejection, the average (*SD*) number of trials retained on Tone trials were 76.57(3.34), 76.43(3.30), and 75.09(4.48) for Position 1, 2 and 3, respectively (~95%). For LAS trials, an average of 17.91(3.16), 19.00(1.57) and 19.00(1.28) trials were retained for each respective Position (~89-95%). For EEG and blink latency analyses, non-response and missing blinks were also excluded resulting in a retained average of 15.48(4.28), 15.7(4.38) and 14.74(4.95) trials for respective position (~75%).

To examine preparation-related changes in the brain, we measured ERP mean amplitude over a 200 ms interval preceding the Tone at the trial-level. Voltages were measured at sites corresponding to motor (Cz), auditory (T7 and T8 average) and visual (Oz) areas.

### Statistical Analysis

Statistical analyses were conducted in R statistics and R Studio using linear mixed models with the ‘lme4’ package (Bates et al., 2015). We presented the results as *F*-values using the ‘anova’ function. For follow-up pairwise contrasts, we computed estimated marginal means using the ‘emmeans’ function from the ‘emmeans’ package (Lenth et al., 2020). We presented the results of these pairwise categorical comparisons as t-ratios (mean difference estimate divided by standard error) and *p*-values for multiple comparisons were corrected using the Hochberg method.

For the finger response and ERP data, we modelled the dependent variable (movement onset RT, peak force, pre-stimulus amplitude) with Position (1,2,3), Stimulus-Type (tone, LAS) and their interaction as fixed-effects. For the eye-blink startle reflex, we analysed the dependent variable (blink onset latency, blink amplitude) on LAS trials with Position (Warning flash, Position 1, 2, 3) as the only fixed-effect. For all models, intercepts for participants and trial history (number of trials since the last LAS) were modelled as nested random effects.

## Results

### Finger Response

#### Response Time

RTs were shorter on LAS trials (*F*_(1,6388)_ = 148.63, *p* < .001**) and decreased as sounds were presented later (*F*(2, 6388) = 951.51, *p* < .001**), reflecting effects of stimulus-intensity and preparation-level. A two-way interaction showed that RT differences decreased as position increased, possibly reflecting a floor effect as reactions approach their lower-limit (*F*_(2,6388)_ = 4.34, *p* = .013*; RT differences across position: 22.3 ms _*p* < *.001*_ **→** 18.0 ms _*p* < *.001*_**→** 12.1 ms _*p* < *.001*_).

According to the classical model, not all responses to a LAS presented at 104 dB are expected to activate the sub-cortical mechanisms responsible for the StartReact effect (Carlsen et al., 2007; but also see Marinovic & Tresilian, 2016; McInnes et al., 2020). It is possible that this mechanism was only engaged on a subset of super-fast LAS responses. As such, effects of motor preparation would be absent on super-fast responses, as this mechanism by-passes voluntary control processes. To investigate this, (1) we compared cumulative distribution functions (CDFs) of RTs between tone and LAS trials and (2) we separately analysed a subset of fast responses. CDFs have been demonstrated to be a more sensitive and reliable way of quantifying RT effects in the context of StartReact compared to traditional methods which only analyse a small subset of trials in which a startle reflex in the sternocleidomastoid muscle are observed (McInnes, Castellote, et al., 2021).

The results of the CDF analysis was consistent with our main analysis, we observed main effects and interactions between Stimulus-Type, Position (stimulus-type, *F*_(1,1298)_ = 243.70, *p* < .001**; position, *F*_(2,1298)_ = 1058.62, *p* < .001**; stimulus-type × position, *F*_(2,1298)_ = 8.89, *p* < .001**). Notably, percentiles did not influence the two-way interaction (stimulus-type × position × percentile, *F*_(18,1298)_ = 0.16, *p* = .999), demonstrating that the effects of stimulus-type and position are evident throughout the entire RT distribution. In our analysis of super-fast responses (5^th^ – 25^th^ percentiles), the same pattern of effects were observed (stimulus-type, *F*_(1,386)_ = 62.67, *p* < .001**; Position, *F*_(2,386)_ = 345.20, *p* < .001**; stimulus-type × position, *F*_(2,368)_ = 5.40, *p* = .001**).

**Figure 4.**
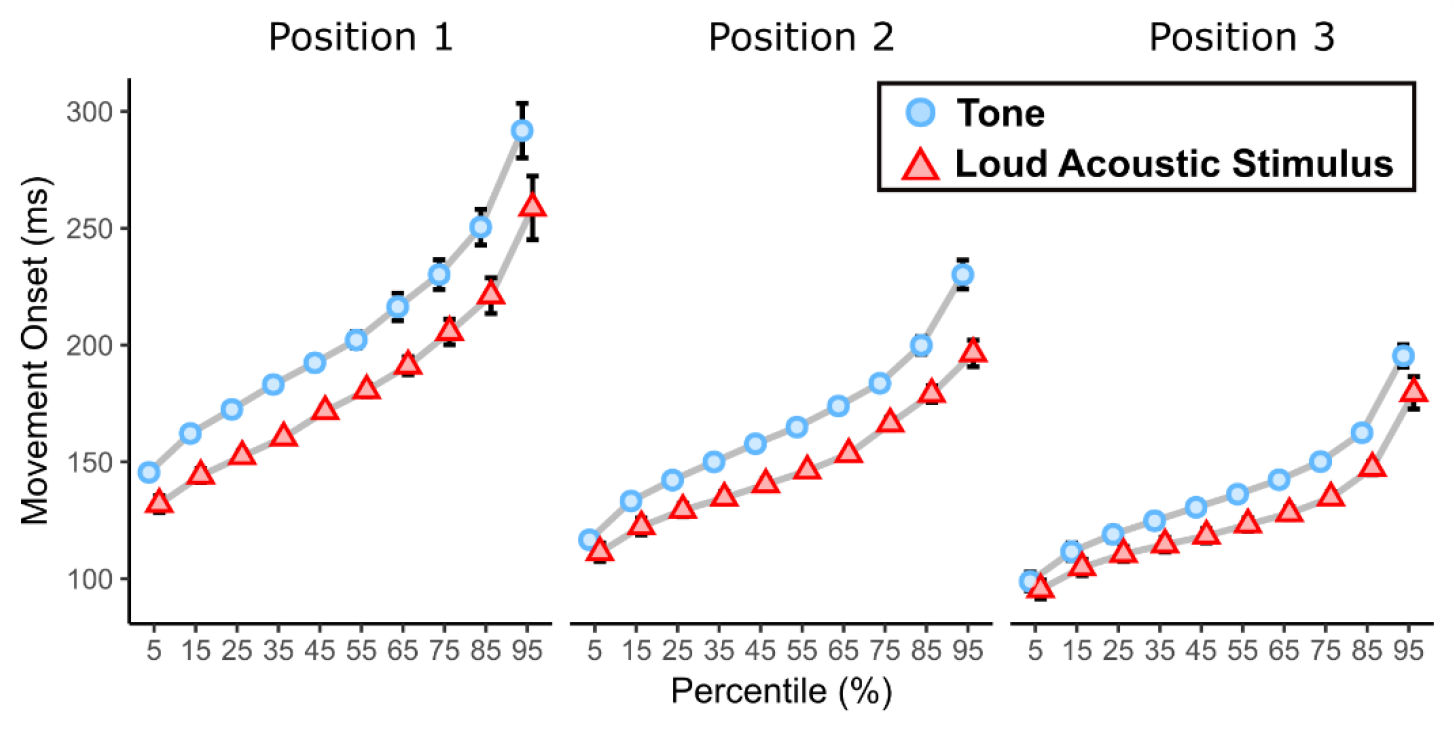
Cumulative distribution functions for RT, showing grand means with within-subject error bars for each percentile (5-95%) and Stimulus-type (tone, LAS) at each Position (1-3).

#### Peak Force

Overall, the force of responses increased as Position increases (*F*_(2,6679)_ = 17.56, *p* < .001). However, peak force was only enhanced on LAS trials (relative to tone trials) at Position 3 (Stimulus-Type × Position, *F*_(2,6679)_ = 5.70, *p* = .003; Force differences: 0.04 N _*ns*_ **→** 0.05 N _*ns*_ **→** 0.34 N _*p* < .001_). Further inspection of the data showed that this pattern was consistent across all blocks.

### Eye-Blink Startle Reflex (LAS trials only)

#### Blink onset latency

We observed a gradual decrease in blink latency as Position increased (*F*_(3,1354)_ = 19.42, *p* < .001). This decrease diminished across position, with the largest difference between the Warning and 1^st^ position, with smaller reductions there-after (Latency differences = 2.72 ms _*p* < .001_ **→** 1.21 ms _*p* = *.051*_ **→** 0.08 ms _*p* = *.868*_). Follow-up analyses showed a positive association between RT and blink latency, where shorter RTs were associated with earlier blink onset latencies (*F*_(1,614)_ = 10.43, *p* < .001, β = 0.60). This effect remained when also accounting for LAS Position. However, there was no association with peak force (*F*_(1,1069)_ = 0.08, *p* = .780, β = 0.00).

#### Blink amplitude

We did not observe an effect of Position for blink amplitude (*F*_(3,1359)_ = 0.99, *p* = .396). To investigate whether habituation may have contributed to the absence of amplitude effects, we examined the first block separately but this did not yield a different result (*F*_(3,418)_ = 0.76, *p* = .514).

### Electrophysiological data

***Motor Area***

Overall, pre-stimulus amplitude at Cz became more negative as Position increased (*F*_(2,6129)_ = 29.49, *p* < .001). However, no significant effect of Stimulus-type (*F*_(2,6129)_ = 0.73, *p* = .392) or two-way interaction were observed (*F*_(2,6129)_ = 0.77, *p* = .461). Follow-up analyses revealed a positive association between Cz activity and RT (*F*_(1,6133)_ = 54.28, *p* < .001, β = 0.11).

#### Auditory Area

Similarly, pre-stimulus amplitude at T7 and T8 also decreased as Position increased (*F*_(2,5927)_ = 16.92, *p* < .001). However, no significant effect of Stimulus-type (*F*_(1,5927)_ = 1.34, *p* = .248) or two-way interaction were observed (*F*_(2,5927)_ = .43, *p* = .648, *R^2^p* < .001). Follow-up analyses revealed a positive association between activity in T7/T8 and RT (*F*_(1,5931)_ = 22.89, *p* < .001, β = 0.11).

#### Visual Area

Unlike effects observed in motor and auditory areas, no statistically significant effects or interactions were observed at Oz (Position, *F*_(2,6127)_ = 2.34, *p* = .097; Stimulus-type, *F*_(1,6217)_ = 1.94, *p* = .164; Interaction, *F*_(2,6217)_ = 0.21, *p* = .812).

## Discussion

In this study, we examined the activation model account of the StartReact effect which argues that the effect is not a special phenomenon mediated by startle-reflex pathways, rather a particular manifestation of intense stimulation when the nervous system is in a high state of preparation. To test the hypothesis that LAS responses can be explained by a combination of preparation state and stimulus intensity, we induced different levels of preparation by altering the conditional probability of the imperative stimulus over the course of each trial. According to the activation model, we predicted that higher preparation-levels would be associated with increased neural activation and reduced response times – demonstrating that motor actions are set-up by preparatory processes which alter the state of the nervous system. We also predicted RTs to LAS would be reduced compared to tones – but would show the same effects of preparation-level on RT and neural activation.

As expected, increasing preparation-level was associated with facilitated movements (reduced RT, increased force) and increased neural activation (increased negativity in motor and auditory scalp areas, and reduced blink onset latencies). With respect to the StartReact effect, shorter RTs were observed on LAS trials, and the same pattern of preparation-level effects on RT and neural activation were present on both tone and LAS trials. Collectively, these results comment on the nature of movement preparation as (1) an evolving process shaped by expectations about when the action is likely to be required, and (2) a distributed process that also engages areas beyond the motor system. The results also illustrate that reactions to LAS can be explained by a combination of established phenomena and the existence of separate sub-cortical pathway is not necessary to explain the StartReact effect – at least for cortically-mediated actions such as individual finger movements (Marinovic & Tresilian, 2016). Although, our data does provide evidence that sub-cortically mediated actions (eye-blink reflexes) can also be modulated by cortical preparatory processes. In the following sections, we discuss how our manipulation of the conditional probability allows for these insights.

### Motor actions are prepared strategically according to the stimulus probability

The effect of the preparation-level manipulation on RT and neural activation demonstrates how motor actions and the underlying processes that set-up the system are shaped by expectations about when a response is likely required. Although there were an equal number of trials for each temporal position (i.e., the global probabilities were the same), RTs decreased with each position. This suggests that participants were updating their expectations based on the absence of tones, gaining more confidence that the tone will occur on the next flash – following the evolving conditional probability of the imperative stimulus represented in Table 1. This RT reduction likely reflects a strategic trade-off between speed and accuracy, as opposed to physiological limits on how quickly actions can be prepared. Limiting the level of preparation at earlier positions may serve to prevent premature and incorrect responses.

This idea is consistent with recent findings by Leow et al. (2018) who isolated the effects of expectancy by using the same flash-based design to contrast predictable and unpredictable (same as ours) variants. In their predictable task, imperative stimuli were also presented at different but known temporal positions. Across the board, RTs were significantly shorter in the predictable compared to the unpredictable task. Notably, mean tone RT at the first position of the predictable task was ~145 ms, which is comparable to our tone RTs at the final position (~140 ms) – demonstrating that the nervous system can reach a high-level of preparedness by the first flash (within 600 ms). Classical experiments studying the lower limits of preparation have shown that optimal RTs can be reached around 200-300 ms after the warning cue (Alegria, 1974; Müller-Gethmann et al., 2003) – about half of the time of the first flash. The fact that the nervous system appears to be under-prepared in early positions when there is temporal uncertainty demonstrates that preparation is not simply about the programming and passive storage of motor commands to be later released, rather a state that must be actively controlled and maintained. The gradual reduction in RT suggest that preparation-states are unsustainable and difficult to precisely control – otherwise we would observe consistent RTs across preparation-levels. The slower RT in early positions likely reflect a balance between speed, accuracy, and effort given the low likelihood of the required response (~29%).

In activation models, preparation is conceptualised described as a continuous but single-stage process where an action is triggered when activation crosses the initiation threshold. However, some researchers have also proposed two-process models which separate preparation and initiation processes (Haith et al., 2016; Weinberg, 2016). Preparation and initiation processes are theorised as parallel-but-staggered processes that specify the ‘what’ and ‘when’ of movements, which have been used to explain why voluntary responses can take ~ 50 – 100 ms longer to initiate compared to reactions to perturbations, how actions may be initiated before preparation is complete, and other phenomenon like self-paced movements. In the context of our study, this framework can offer two alternate interpretations: Firstly, given that our task uses a very simple action (choiceless finger press) which may not require preparation, differences in RT across preparation-level may reflect a modulation of the initiation process as opposed to the preparation. Secondly, reactions to LAS may be faster because the stimulus-intensity effect is specifically speeding-up the initiation process. Although the distinction between preparation and initiation does not meaningfully change our interpretation that preparation is associated with evolving and wide-spread changes in brain dynamics, the distinction between processes may be important in future studies considering more complex movements and high-urgency situations – when actions might be initiated before movement preparation is complete.

Further commenting on the strategic nature of movement preparation, we observed a peculiar yet consistent reduction in peak force on the final position on tone trials. This reduction might reflect the engagement of inhibitory mechanisms to avoid false-starts in the event of a Catch trial. Interestingly, this reduction was not observed for LAS trials, rather we observed the greatest increase in peak force on LAS trials at the final position. The pattern of force results is difficult to interpret as this enhancement was specific to the final position, but one possibility is that response force might only be facilitated by intense stimuli when preparation-levels are relatively high. Although this effect remains to be further examined, it highlights the potential impact of Catch trials on task strategy and therefore movement preparation.

### Activity in motor and auditory areas reveal the distributed nature of movement preparation

The coupling between RT and pre-stimulus cortical activation at the condition- and trial-levels are consistent with the notion that preparation is marked by changes in both the excitability of the nervous system and the motor action. The evolving nature of this process is evident in the ERP waveform, where step-like changes can be seen as expectations about the likelihood of a response are updated with each flash (See Figure 2). While preparation-level effects show the shorter timescale predictions about the likelihood of a response, trial-level associations between RT and cortical-activation are likely reflect longer timescale (across-trial) predictions about the occurrence of the imperative stimulus.

Interestingly, preparation-level effects on cortical activation were evident in motor as well as auditory scalp regions, demonstrating that preparation is a distributed process. This suggests that the ability to react is not only dependent on the state of the motor system, but also the state of sensory areas which could allow for earlier detection the imperative stimulus leading to earlier responses. This is in line with recent work by Li et al. (2017) showing activation of the auditory cortex is associated with the initiation of motor actions in mice.

Given that our task also had a visual component, we also hypothesised a preparation-related increase in activation at visual areas – in-line with previous work by Bueti and Macaluso (2010) who found that auditory expectations also modulated activity in visual areas during movement preparation. However, this effect was not reliable, which may be due to the lack of task-relevant information provided by the visual flashes (i.e., flashes only provided generic but not specific information about the appearance of imperative stimuli). Alternatively, the absence may be related to the specific use of sounds which may elicit multi-sensory representations. For example, Bueti and Macaluso (2010) used sounds such as the those of hand-clapping or of a hammer-hammering which can be visually imagined, but we used pure tones and broad-band white noise which are not naturally associated with such visual imagery (but see Swallow et al., 2012). Overall, the presence of preparation-level effects in motor and auditory, but not the visual area demonstrates that while movement preparation is a distributed process, it appears to selectively engage only task-relevant areas.

### Movement preparation also influences the excitability of sub-cortical circuits

Lastly, our task provides new insights regarding the effects of movement preparation on startle-related circuits. To date, numerous studies have used LAS as a probe to study the time-course of changes in sub-cortical excitability during movement preparation by delivering the LAS at different times: before, with, or after the imperative stimulus. Collectively, there is evidence for significant modulation of the eye-blink reflex shortly before and after the presentation of the imperative stimulus in reaction-based tasks (e.g., Lipp et al., 2007; Lipp et al., 2001; Marinovic et al., 2013). In anticipatory timing tasks, modulation of the eye-blink reflex was able to detect a phenomenon known as pre-movement inhibition (e.g., McInnes, Lipp, et al., 2021; Nguyen et al., 2021). However, the specific time-course and direction of these effects do vary, with some studies reporting null effects for the eye-blink reflex (Kumru et al., 2006; Kumru & Valls-Solé, 2006). A major difficulty with interpreting these discrepant findings is that response requirements and contextual parameters can vary significantly across studies which can have dramatic impacts on the time-course of preparation (e.g., choice response vs. single response, jittered vs. fixed inter-stimulus intervals, equiprobable vs. skewed stimulus/response probabilities, and presence of catch trials).

Our current design offers a different approach to studying movement preparation by allowing us to systematically manipulate the amount and the time-course of preparation, as opposed to standardising the presentation of trials (in fixed-cue RT and anticipatory tasks). Using this design, we were able to show that changes in sub-cortical excitability occur alongside changes in the motor-response and cortical activation during preparation. In classical models of the StartReact effect cortical and sub-cortical circuits are given different roles – where sub-cortical circuits only become relevant after preparation is ‘complete’ and the resultant motor programme is transferred sub-cortically for storage and triggering (Valls-Solé et al., 1999). However, our data demonstrates that changes in sub-cortical excitability are part of the entire preparation process – possibly serving to facilitate the transmission of the motor action.

Although blink latency was associated with preparation-level, blink amplitude did not. This discrepancy may be attributed to onset latency and peak amplitude measures capturing different times in the EMG signal. It is known that intense stimuli can elicit two distinct eye-blink components: the auditory eye-blink reflex, and the auditory startle reflex. The auditory eye-blink reflex occurs at short latencies and is thought to be mediated by mesencephalic circuits, and the auditory startle reflex triggers a later response along with a generalised skeletomuscular response – thought to originate from bulbopontine circuits, distinct form those associated the auditory eye-blink (Brown et al., 1991). Given that blink latency captures the onset of EMG activity, it is likely to capture the auditory eye-blink reflex whereas the peak amplitude is more likely to capture the auditory startle reflex (if larger). Although these measures may reflect activity of separate circuits, there is some overlap. In Nguyen et al. (2021) we were able show evidence of eye-blink suppression in both amplitude and latency. In addition to discrepancies caused by task-differences, not all studies report both blink amplitude and latency which makes it difficult to evaluate the two metrics. Nevertheless, our data provides positive evidence that the excitability of sub-cortical startle circuits are modulated by the level of preparation.

## Conclusion

The results of this study demonstrate that responses to LAS can be explained by a combination of multisite (e.g., motor, and auditory) preparation states and stimulus intensity. RT and neural activation evolved with the increasing conditional probabilities of the imperative stimulus, suggesting that preparation was based on the updating expectations occurring throughout the course of each trial – reflecting a strategic optimisation between speed and accuracy. As predicted by the activation model, preparation effects were evident on both LAS and tones. Our task design provides a useful method for systematically manipulating movement preparation which allows us to show its evolving and widespread (but selective changes) effects on the nervous system.

## Acknowledgements

The study was supported by a Discovery Project grant from the Australian Research Council (DP180100394) awarded to W.M. and O.V.L.

## Data Accessibility Statement

The summarised data and analysis code will be made publicly available on GitHub once the manuscript has been peer-reviewed and published (https://github.com/angnguyen)

